# Photon count estimation in single-molecule localization microscopy

**DOI:** 10.1101/396424

**Authors:** Rasmus Ø. Thorsen, Christiaan N. Hulleman, Mathias Hammer, David Grünwald, Sjoerd Stallinga, Rieger Bernd

## Abstract

Recently, Franke, Sauer and van de Linde^1^ introduced a way to estimate the axial position of single-molecules (TRABI). To this end, they compared the detected photon count from a temporal radial-aperture-based intensity estimation to the estimated count from Gaussian point-spread function (PSF) fitting to the data. Empirically they found this photometric ratio to be around 0.7-0.8 close to focus and decreasing away from it. Here, we explain this reported but unexplained discrepancy and furthermore show that the photometric ratio as indicator for axial position is susceptible even to typical optical aberrations.

---

In Fig. 1A we show the photon count from a 45 nm bead imaged with an aberration-corrected microscope^2^ (see **Supplementary Methods** for details) estimated by three different methods (Gaussian PSF fit, TRABI, Vectorial PSF fit^3^) as a function of aperture radius or fit box size, respectively (for reproducibility see Supplementary Fig. 1). It is evident that the estimated count increases with increasing area for all three methods, i.e. no method finds the true count for a realistic area as the true microscope PSF has a very long tail.Simulations of full-vectorial PSFs support this conclusion (Supplementary Fig. 2), showing that the tail deviates substantially from the Airy PSF model^3^. It is also evident that with any aperture based method the true count can only be approximated up to 90% with aperture radii less than one micron (Supplementary Fig. 3) and that Gaussian PSF fitting performs worse as a Gaussian cannot fit the long tail at all. This, however, does not bias the localization estimate of Gaussian fitting for round spots^3^. The suitability of sub-diffraction sized beads for these experiments was investigated in simulation and found to increase the FWHM by only a few nanometers compared to the single-molecule PSF (Supplementary Fig. 4) while giving access to more light over a longer period during the experiment.

**Figure 1.**
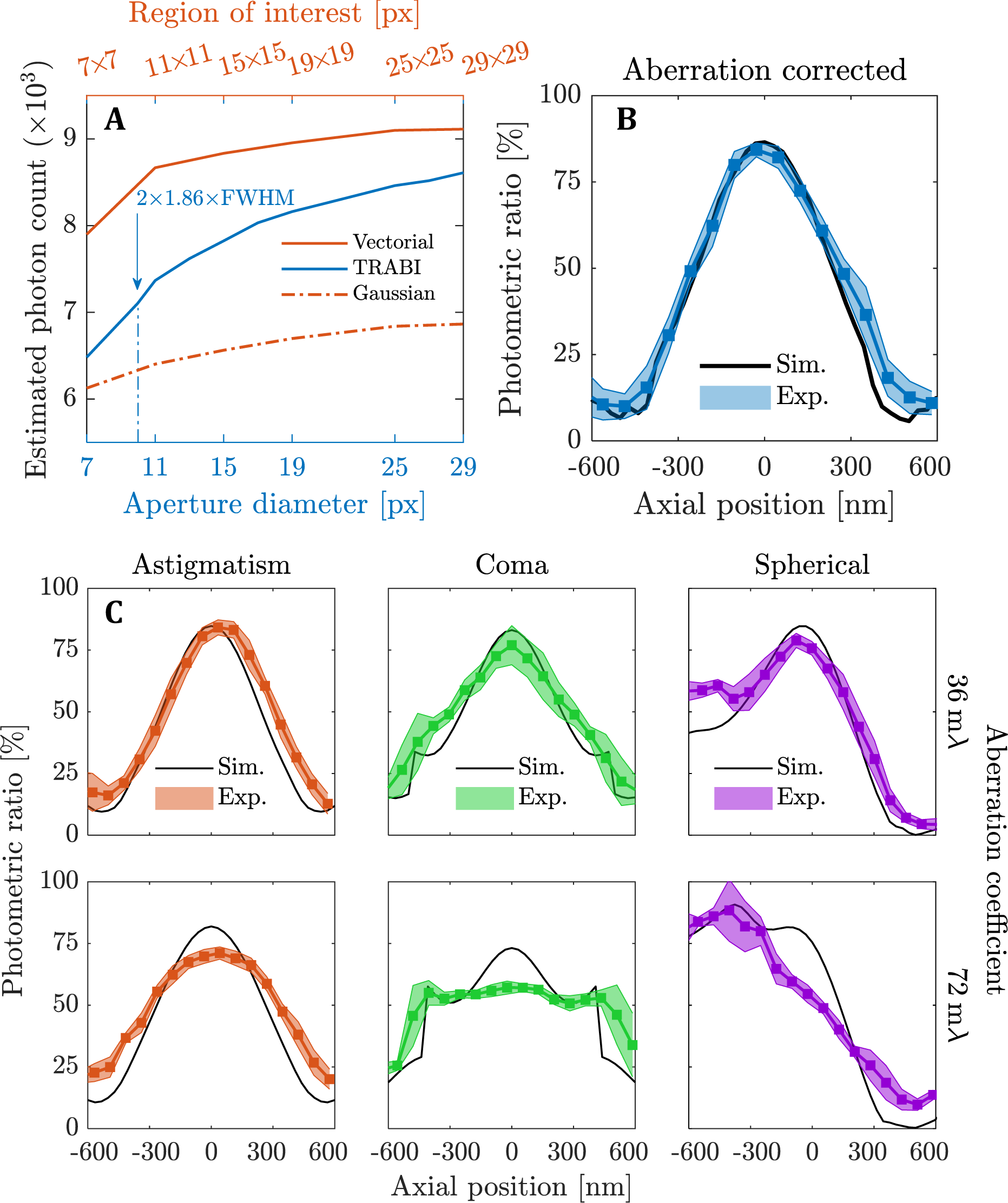
Photon count estimation and the effects of aberrations on the photometric ratio. **(A)** Estimated photon count for a 45 nm diameter bead imaged with an aberration corrected microscope as a function of analysis area on the camera. Three lines show the count for fitting with a fully-fledged vectorial or simplified Gaussian PSF model compared to TRABI, FWHM= 214.5 nm; pixel size= 80 nm. Aperture diameter of 2×1.86×FWHM as suggested by Franke et al.^1^. **(B)** The photometric ratio (Gaussian fit over TRABI value) over six bead measurements as a function of the axial position. The colored error band indicates one standard deviation. Area of fit 7×7 pixels and aperture radius 1.86×FWHM. **(C)** Effect on the photometric ratio for beads single-mode aberrated with root-mean-square aberration coefficients set at half (36 mλ; first row) and full diffraction limit (72 mλ; second row).

Next, we varied the axial position of the sample while imaging aberration corrected beads and evaluated the photometric ratio between photon count estimates from Gaussian fitting and TRABI as a function of defocus, as shown in Fig. 1B (see Supplementary Fig. 2 for sensitivity to fit area). The residual wavefront aberration was 24 mλ RMS (see Supplementary Fig. 5 for experimentally retrieved aberration coefficients). Simulations using the fitted residual aberrations result in photometric ratios that agree well with experiment. We find a photometric ratio of 85% in contrast to the values around 75% in focus reported by Franke et al.^1^, which we attribute to aberrations present in their experiment. To assess the influence of aberrations, we experimentally engineered PSFs with small amounts of astigmatism, coma or spherical aberration. Photometric ratios obtained from these experiments match those obtained from simulations with added aberrations (see Fig. 1C). The maximum value of the photometric ratio in focus, overall shape and values strongly depend on the aberrations, resulting in curves that are either broadened, flattened or made asymmetrical. The amounts of added aberrations still represent a lens that sells as diffraction limited (Maréchal diffraction limit is at 72 mλ), indicating that these aberration levels and combinations thereof are seen in typical setups. We estimated the impact of these small aberrations on the expected axial position error by comparing against an aberration corrected calibration and find errors between ±100 to ±200 nm over 800 nm dynamic range (see Supplementary Fig. 6). We inspected seven different setups for aberrations and found that typical non-corrected systems can have axial errors on the order of ±50 to ±100 nm (see Supplementary Fig. 7). Sample induced refractive index mismatch, e.g. by using oil immersion into a watery enviroment, leads to spherical aberration but also non-spherical components^4^ on the same order as we simulated here. We conclude that in order to convert the photometric ratio to a viable, accurate depth map the optical aberrations must be known to a very high degree (wave front uncertainty < 10 mλ results in axial uncertainty < 20 nm).

## Data availability

The data is available for download at https://data.4tu.nl/download/uuid:ea2ea179-26f4-4e1a-90e1-b2759b553ce8/. The software is available as Matlab scripts in open-source from ftp://qiftp.tudelft.nl/rieger/outgoing/Rasmus_photoncount.zip

## Acknowledgments

B.R., C.N.H. acknowledge European Research Council grant no. 648580 and B.R., R.Ø.T., D.G. acknowledge National Institute of Health grant no. U01EB021238. We thank Job Dekker for providing access to serval microscopes and Keith Lidke for providing PSF data.

## Author contribution

RØT performed simulations and analyzed data, CNH performed experiments, MH and DG provided 3D PSF data from several microscopes, SS and BR designed and coordinated the research. BR, SS, RØT wrote the manuscript, and all authors commented on it.

## Supplementary Methods

### Optical microscope setup

The experimental data is acquired with a setup that consists of a Nikon Ti-E microscope with a spatial light modulator (SLM) for aberration correction and PSF engineering, as described by us earlier^1^. In short, beads are excited by a 488 nm laser (Sapphire 488-100 CW CDRH, Coherent) through a dichroic filter set (Ex: Semrock FF01-460/60-25, Di: Semrock Di02-R532-25X36, Em: Semrock FF01-545/55-25) and an objective lens (APO TIRF 100x/1.49, Nikon). The emission fluorescence passes (peak λ = 552 nm) through a relay system (f1 = 100 mm, Thorlabs AC254-100-A and f2 = 200 mm, Thorlabs AC508-200-A) resulting in a final magnification of 200x. The SLM (XY-series, 512×512, 15 μm pixel size, Meadowlark) is placed in the Fourier plane of the relay system. The fluorescence image signal is captured on an EMCCD camera (iXon Ultra - X987, 512×512, 16 μm pixel size, Andor) which has in effect a back-projected pixel size of 80 nm in the object plane.

### Sample preparation

Fluorescent beads of 45±8 nm diameter (FluoSpheres 0.04 μm 505/515, ThermoFisher) with peak emission at 552 nm through our filter set are used in the imaging experiments. The beads are immersed in a mounting medium of immersion oil (n=1.518 Type F, Nikon) to match the refractive index to the coverslip and immersion oil used for the objective lens. The beads are diluted to 10^11^ particles/mL in water, and 2.5 μL of the solution is drop-cast on a coverslip. After evaporation of the water, a drop of immersion oil is applied, and the coverslip is glued around the edges with nail varnish to a microscope slide. This step ensured the rigidity of the sample and is necessary for precise z-positioning.

### Data acquisition

Fluorescent beads are imaged with 3.5 mW of excitation power (measured at the back aperture of the objective) and an exposure time of 1 second. For each set of aberration (type and magnitude), the experiment is repeated three times in the same field of view (FOV) to test for reproducibility. For each FOV, we select two beads located in regions with a low bead density to avoid crosstalk of beads. This is done to ensure a large region of interest with the no/minimal overlap of their respective PSF shoulders. This results in a total of six configurations, i.e. two beads measured three times. These six configurations are used to compute the mean and standard deviation of the data as shown as error bars in Figure 1B,C and Supplementary Figure 5. The data in Figure 1A and Supplementary Figure 1 and **2B,C** are for individual beads.

### Aberration retrieval and correction

Our setup contains an additional light path for pixel-wise calibration of the LCoS SLM to ≤20 mλ RMS wavefront aberration as described earlier^1^. After calibration of the SLM, we acquire a z-stack of the fluorescent bead by moving the piezo stage in steps of 80 nm from −800 nm to +800 nm around focus. The z-stack is fitted with a 3D full vectorial PSF model in order to estimate the aberrations of the optical system^1^. The retrieved aberration coefficients are fed to the SLM with opposite sign in order to correct for the aberrations. Subsequently, another z-stack is acquired and fitted to verify the compensation of the aberrations. We then add aberrations of a desired type and magnitude deliberately in order to investigate the impact of aberrations in a well-controlled fashion.

### Simulation of a realistic PSF model

We use the vectorial PSF model also for simulating the effect of arbitrary aberrations on photon counting^7^. In addition we take into account the non-zero size of a bead by convolving the PSF with the bead size. This fully-fledged PSF model takes the following effects into account: interfaces between media, polarization, dipole orientation (here freely rotating dipoles are assumed) and type and magnitude of aberrations. The simulation parameters are based on our optical setup: i.e., medium refractive index n= 1.518, numerical aperture NA= 1.49, wavelength λ= 552 nm, backprojected pixel size = 80 nm. We assume that the refractive indices between the imaging medium, coverslip, and immersion medium are matched (based on the immersion of the beads in oil). In Supplementary Figure 3 and **4** we use a smaller sampling distance of 1 nm to exclude quantization effects. The bead diameter is 45 nm except in Supplementary Figure 4 where the bead diameter is varied from 0 to 180 nm. Results for the Airy PSF model shown in Supplementary Figure 3 follow directly from the derivation as detailed in Born & Wolf^4^ (Chapter 8). The simulated PSFs are normalized such that the sum over the detection plane is unity, then multiplied by the desired number of signal photons and a constant number of background photons per pixel is added before applying shot noise.

### Photon count estimation

To determine the gain and offset, the EMCCD camera is calibrated using the procedure described by van Vliet et al.^2^. For these experiments, we used an EM gain setting of 50 to reduce the read-noise to <1 e-RMS while retaining a good dynamic range and linear response (maximum intensity is kept under 5000 ADU - the linear regime for our camera settings). Excess noise in EMCCD cameras leads to an overestimation of the gain by a factor of two^3^, but this does not pose any problems in the further analysis of the data as only ratios of photon count estimates are of interest here.

The fitting procedure with a vectorial or Gaussian PSF model is implemented with a Levenberg-Marquardt iterative scheme for Maximum-Likelihood estimation (MLE) as described in Smith et al.^6^. The number of iterations is terminated by the tolerance in the residual. The tolerance limit is set to 10^−6^ and 10^−4^ for the vectorial and Gaussian fitter, respectively (maximum number of iterations is 75). Typically the number of iterations is less than 20.

TRABI^5^ is designed to take advantage of fluorescent on-off blinking of photoswitchable or photoactivatable fluorophores. TRABI can estimate both signal and background with a “single aperture” by incorporating time information, i.e. estimating the background in the off-state and the foreground signal in the on state. Here we image non-blinking fluorescent beads, which does not allow us to apply the method directly. Instead, we estimate the number of background photons by an aperture located far away from the bead. For background averaging we use 7 successive frames in the through-focus stack as suggested by Franke et al^5^.

### Estimation of error axial position estimation

The error in the axial position estimate from the photometric ratio is done in the following way. The experiments on different beads give the photometric ratio *PR*(*z*) as a function of axial position *z* with uncertainty Δ*PR*(*z*) (Fig. 1B,C). The experimental aberration-corrected calibration curves *PR*(*z*) and *PR*(*z*) ± Δ*PR*(*z*) (Supplementary Fig. 6A) are up-sampled to 1 nm axial steps and stored in a look-up-table (LUT). Given a measured value *PR*_*exp*_, the LUT will give the estimated axial position *z*_*exp*_ with uncertainty Δ*z*_*exp*_. This uncertainty represents statistical variations in the photometric ratio curves due to e.g. variations across the field-of-view of the microscope, and is on the order 25 nm away from focus to 65 nm close to focus (Supplementary Fig. 6B). The estimated axial position change in case aberrations are present as the photometric ratio curves change (Supplementary Fig. 6C). If calibration is done on an aberration-corrected setup, the application of the aberration-free LUT on photometric ratio measurements on aberrated spots will also lead to a bias in the axial position estimate (Supplementary Fig. 6). These inaccuracies can amount up to 100 nm for aberration levels at 50% of the diffraction limit.

### Microscope specifications for PSF retrieval of Supplement Figure 7

**Dataset 1** is obtained with our setup described in **Supplementary Methods** prior to aberration correction.

**Dataset 2** is acquired on a Leica DMi8 microscope. The sample (TetraSpeck Fluorescent Microspheres, 200 nm, well 4, RI: mounting medium 1.472) is excited by a Leica PL 6000, diode laser through a filter set (Chroma ZT/594rpc, Chroma 89100bs), and an objective lens (Leica Plan-Apochromat 63x/1.40 Oil DIC, RI: embedding medium 1.518). The fluorescence image signal (emission peak at λ= 580 nm) is captured on a sCMOS camera (Hamamatsu C11440 ORCA-flash 4.0, detector 13.312×1312mm^2^, pixel size 6.5×6.5 μm^2^, chip size 2048×2048).

**Datasets 3,4,5,6** are acquired on a Nikon Eclipse Ti-E microscope. The sample (TetraSpeck Fluorescent Microspheres, 200 nm, well 4, RI: mounting medium 1.472) is excited by a Lumencor SPECTRA X, diode laser through a filter set (**dataset 3,5:** Ex.: SPECTRA X Chroma 470/24, Em.: SPECTRA X, ET515/30m Single Bandpass, and **dataset 4,6:** Ex.: SPECTRA X Chroma Excitation Filter 550/15, Em.: SPECTRA X, ET595/40m Single Bandpass) and an objective lens (**dataset 3,4:** CF160 TIRF Apo 60x/oil DIC 1.49 NA, **dataset 5,6:** CF160 SR/TIRF Apo 100x/oil 1.49 NA, RI: embedding medium 1.518). The fluorescence image signal (emission peak **dataset 3,5**: λ = 515 nm, **dataset 4**: λ = 570 nm, **dataset 6**: λ = 580 nm) is captured on a sCMOS camera (Andor Zyla 5.5 sCMOS, detector 16.6×14 mm, pixel size 6.5×6.5 μm^2^, chip size 2560×2160).

**Dataset 7** is acquired on an Olympus IX71 microscope. The sample (FluoSpheres F-8789, dark red, Invitrogen, 40 nm) is excited by a HL63133DG, Thorlabs, 637 nm diode laser through a filter (FF01-692/40-25, Semrock), dichroic mirror (650 nm, Semrock) and an objective lens (UAPON 150XOTIRF, Olympus America Inc., NA 1.45, RI: immersion medium 1.52). The fluorescence image signal (emission peak λ = 690 nm) is captured via a 2X magnification relay optics on an EMCCD camera (iXon 897, Andor, Andor Technologies PLC., CCD size 515×515, pixel size 12 μm). Setup details and sample preparation is described further by Liu et al.^8^.

---

**Supplementary Figure 1.**
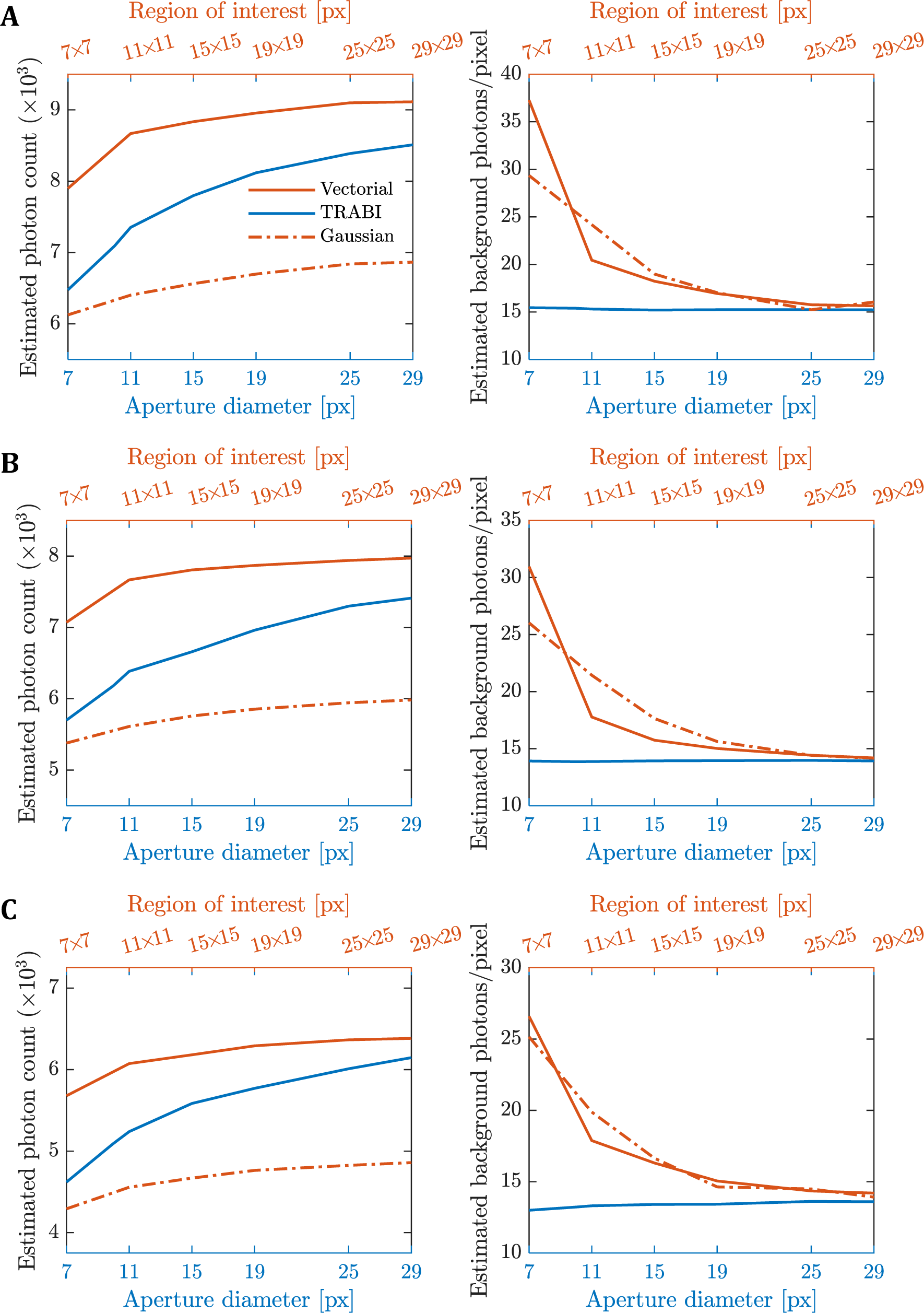
Reproducibility of photon count estimation. **(A-C)** Estimated photon counts and background photons per pixel for three separate 45 nm diameter beads imaged with an aberration corrected microscope as a function of analysis area on the camera, pixel size= 80 nm. The three lines show the count for fitting with a fully-fledged vectorial, TRABI or a simple Gaussian PSF.

**Supplementary Figure 2.**
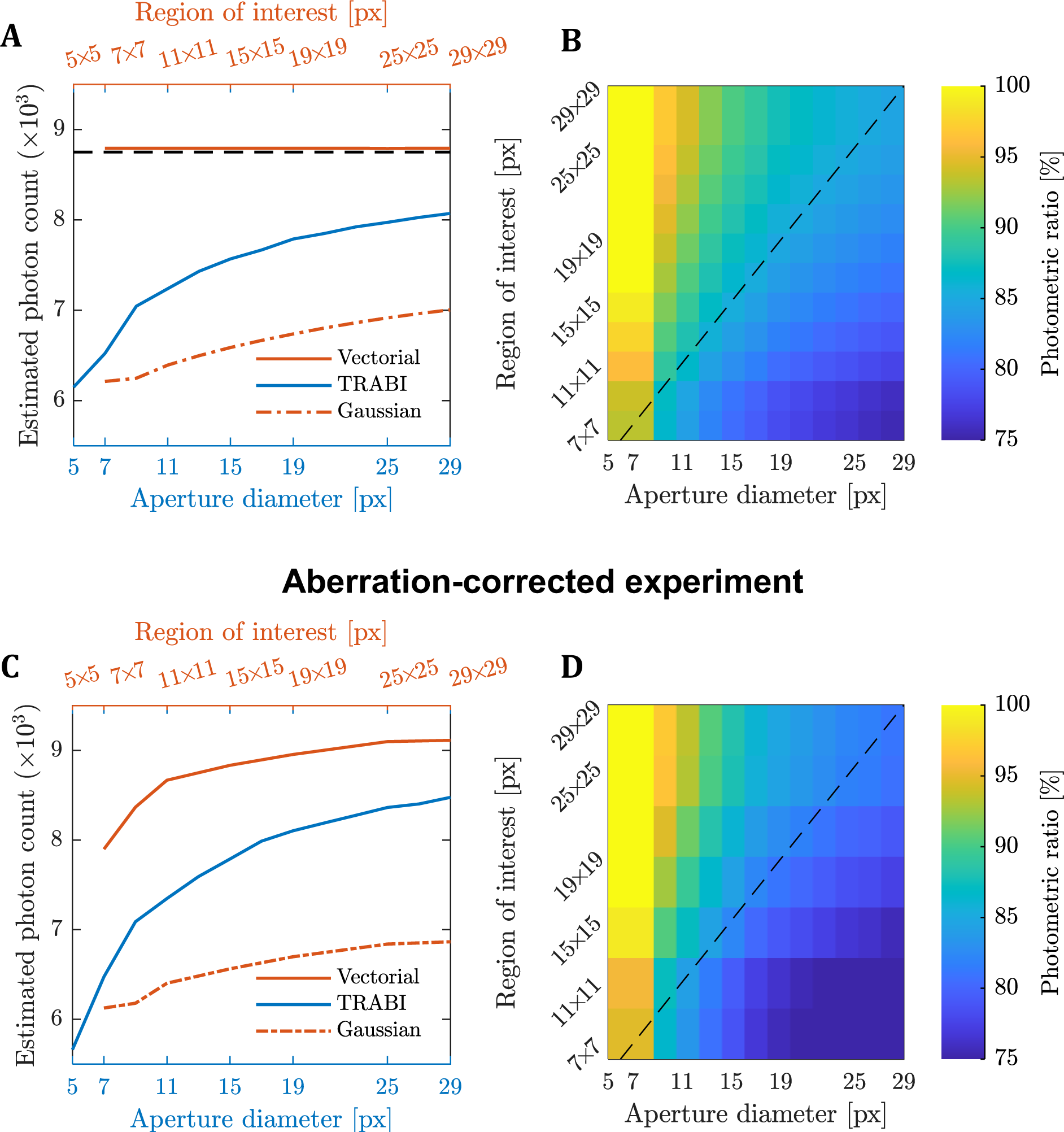
Comparison of photon count estimations in simulation and experiment. **(A)** Photon count estimation for a single simulated 45 nm bead PSF as a function of PSF image area on the camera for three different estimation algorithms: vectorial fit, TRABI, and Gaussian fit (8750 signal photons, 15 background photons, same pixel size, NA and refractive index values as experiment). **(B)** The photometric ratio between Gaussian fit and TRABI photon count estimation as a function of PSF fit size. The dashed line indicates when the circular aperture size fits precisely within the square region of interest. **(C-D)** Same as panels (A,B) but for an experimentally recorded aberration-corrected 45 nm bead, indicating good quantitative agreement between experiment and simulation.

**Supplementary Figure 3.**
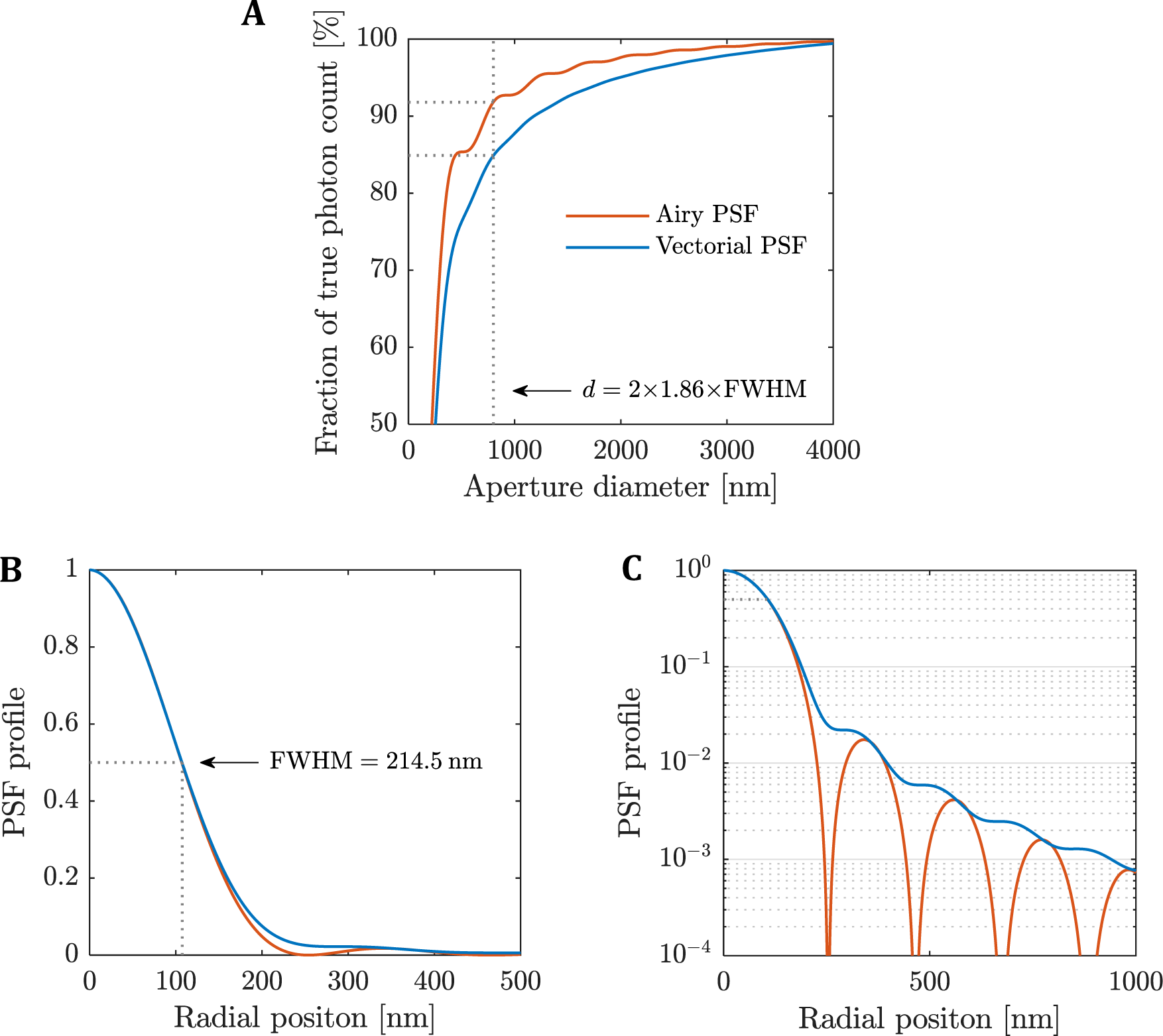
TRABI photon count estimate as a function of aperture size and PSF model. **(A)** Ratio of the estimated photon count by TRABI and simulated number of signal photons captured at the camera as a function of aperture diameter for two PSF models. TRABI aperture diameter (d) of 2×1.86×FWHM (FWHM = 214.5 nm) as suggested in Franke et al.^5^ is indicated. For this aperture size, we find TRABI to estimate 92% and 85% of the total number of photons according to the low NA scalar Airy PSF model and the high NA vectorial PSF model, respectively. The fraction of the total energy contained within an aperture of the prescribed radii in the Airy PSF model (PSF_./01_(ρ) = [2*J*_7_(*D*)/*D*];^2^ where *D* = 2πρ(NA/λ); FWHM = 0.514(λ/NA)) follows the analytically derived formula by Born & Wolf^4^ (Chapter 8; Fig. 8.13), in contrast to the 100% reported by Franke et al.^5^ in their supplement. The PSF simulations were performed with 2500 signal photons and no background photons and a pixel size of 1 nm (to exclude quantization effects). Otherwise, the simulation parameters were the same as the experimental parameters. **(B-C)** Characterisation of the Airy and vectorial PSFs as a function of the radial position in a linear and log scale, respectively, showing the long tail of the PSF.

**Supplementary Figure 4.**
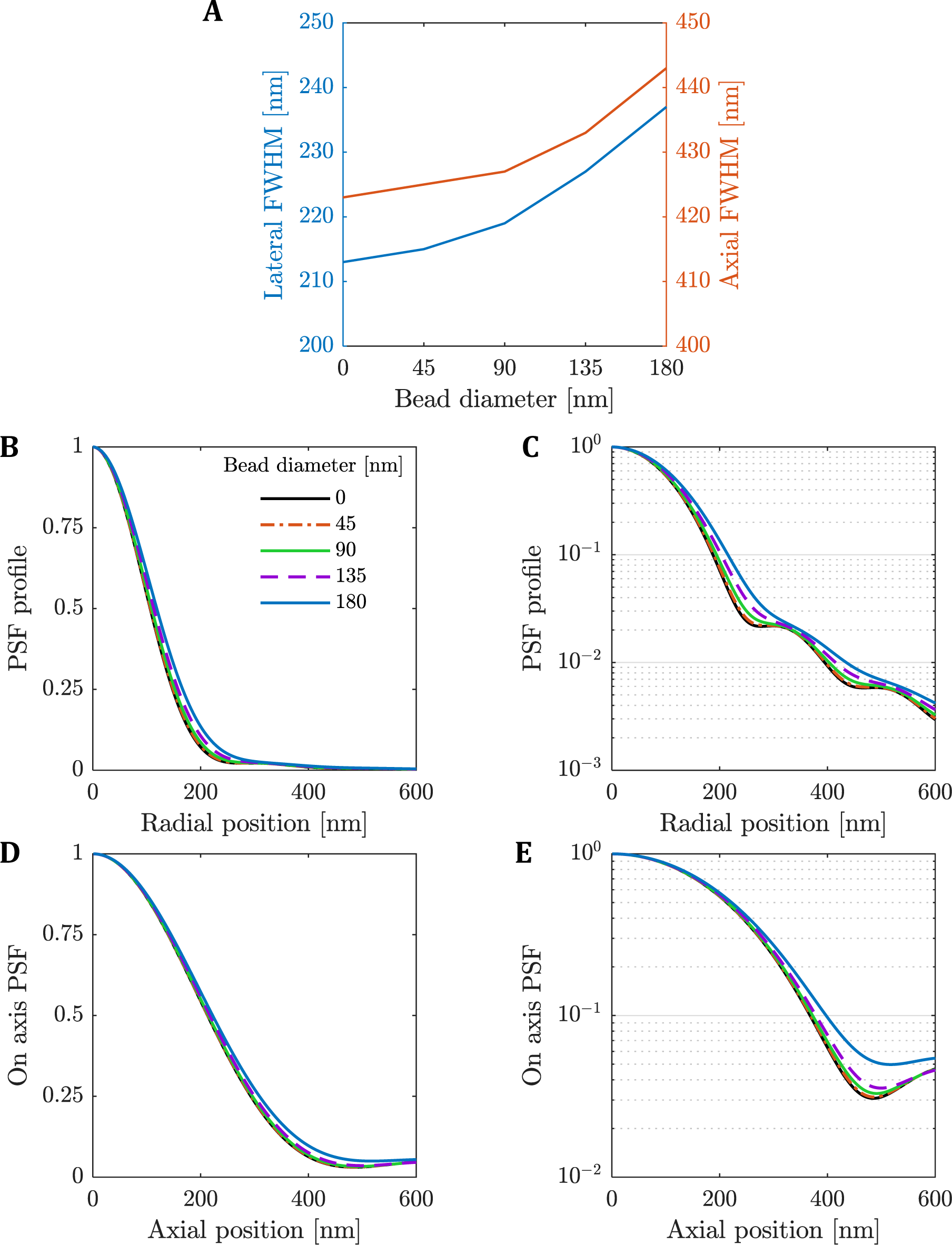
The effect of bead size on the PSF shape. **(A)** The lateral and axial FWHM as a function of bead size. **(B-C)** Lateral average PSF in linear and log scale, respectively, as a function of the radial positon. The PSFs are displayed for different bead sizes. **(D-E)** On axis PSF in linear and log scale, respectively, as a function of the axial position. The 45 nm bead captures the full tail-behaviour in both the lateral and axial direction of an actual single-molecule emitter, and has a FWHM within a few nanometers of a single-molecule emitter, whereas PSF details clearly get lost with larger (>90 nm) beads.

**Supplementary Figure 5.**
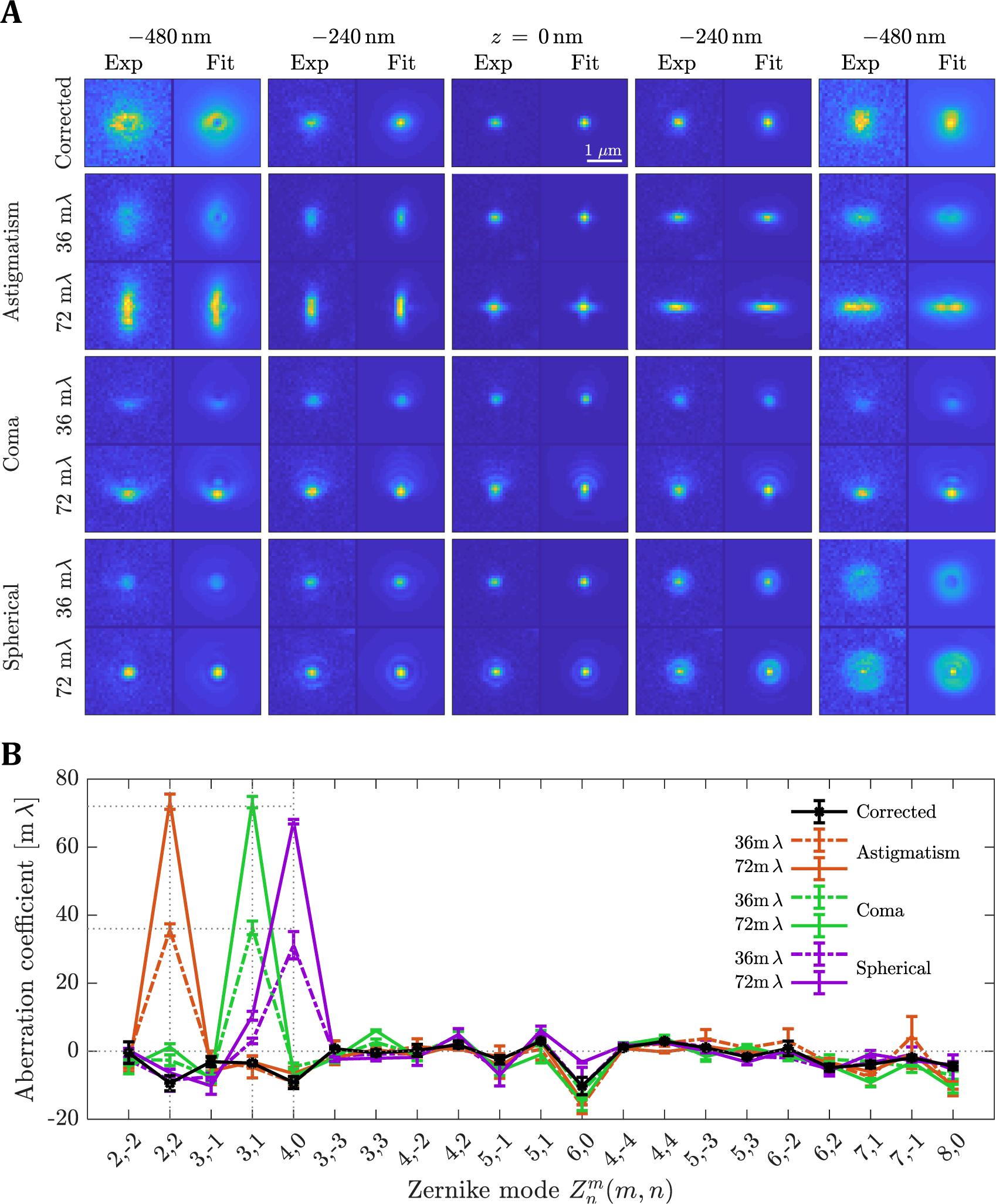
Quantification of aberration retrieval and correction. **(A)** Through-focus PSF image stacks of experimental and fitted PSFs after aberration correction and, subsequently, aberrated with a single primary Zernike mode: astigmatism 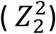, coma 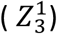, and spherical aberration 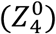, with aberration coefficients 36 mλ and 72 mλ (root-mean-square values). The region of interest for each PSF image is 31×31 pixels with a pixel size of 80 nm. All 4×4 sub-image pairs are contrast stretched with the same factor for better visibility of spot shape. The estimated photon counts were within 9,500-21,000 signal photons and 15-18 background photons per pixel. **(B)** Fitted Zernike modes and retrieved aberration coefficients. The coefficients are averaged over six measurements with error bars indicating one standard deviation. The aberration fit routine includes all tertiary Zernike modes (all 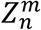 with 2<n+|m|≤8) and assumes optical parameters as described in **Supplementary Methods**. (Horizontal dashed lines are used to guide the eye).

**Supplementary Figure 6.**
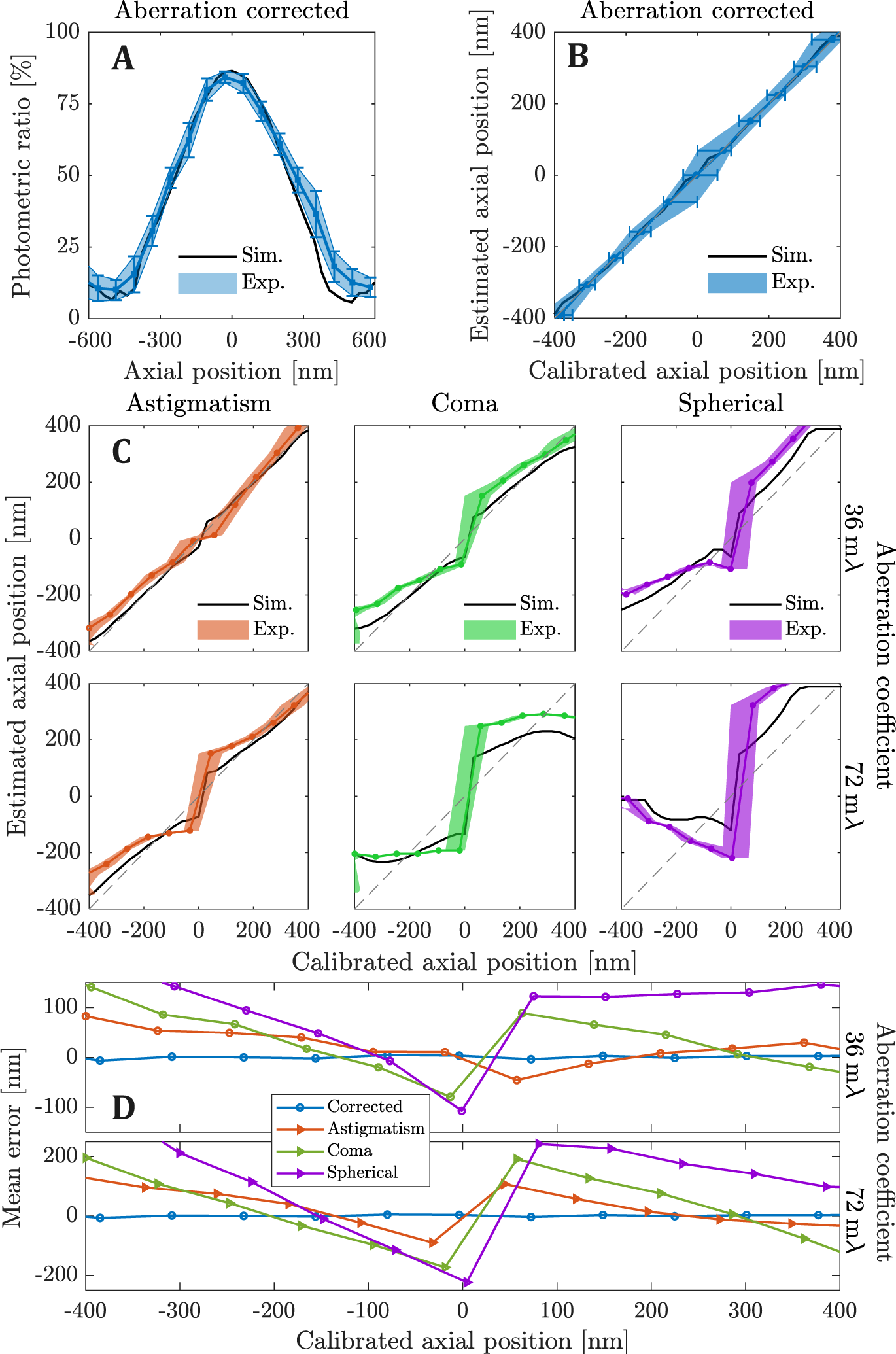
Axial calibration of the photometric ratio and axial estimation error caused by small single-mode aberrations. **(A)** The photometric ratio over six 45 nm bead measurements after aberration-correction (Exp.) and simulated vectorial PSFs (Sim.) as a function of axial position (see **Supplementary Methods**). The error bars indicate one standard deviation. **(B)** The estimated axial position of aberration-corrected beads as a function of the calibrated axial position shown in **A)**. The error bars indicate one standard deviation over the six bead measurements (see **Supplementary Methods** for a description of the error analysis). Around focus the error is a bit larger as expected due to near parabolic shape of the photometric ratio curve. **(C)** Estimated axial position for single-mode aberrations on 45 nm beads using the calibrated axial position from the aberration corrected data in **A)**. The grey dashed line is used to guide the eye for the correct estimation. **(D)** Error of the estimated axial position as the distance between the mean estimated position and the true (calibrated) position.

**Supplementary Figure 7.**
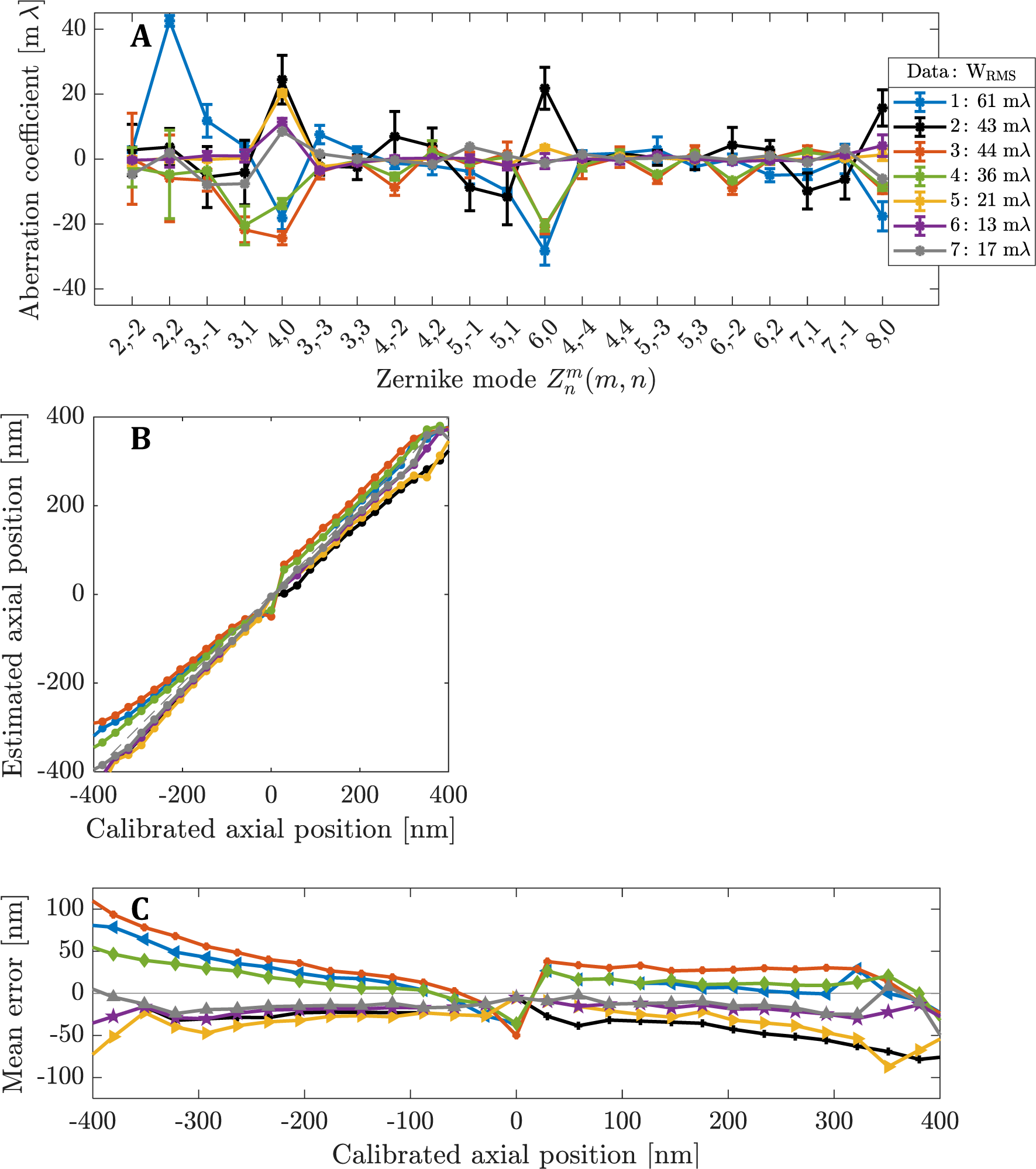
Typical microscope aberrations and their influence on the estimated axial position. **(A)** Fitted Zernike modes and retrieved aberration coefficients for several microscopes, Data 1-7 (specifications are given in **Supplementary Methods**). The coefficients are averaged over six and one bead measurement(s) for datasets 1-6 and 7, respectively, with error bars indicating one standard deviation and W_RMS_ is the mean of the wavefront error. The aberration fit routine includes all tertiary Zernike modes and assumes optical parameters for each microscope as described in **Supplementary Methods**. **(B)** Estimated axial position for simulated single-molecule PSFs with aberrations equalling the mean experimentally found microscope aberrations in **(A)** compared to the calibrated axial position using the aberration-corrected photometric ratio (see Supplementary Fig. 6a). Area of fit 7×7 pixels and aperture radius 1.86×FWHM. **(C)** Error of the estimated axial position taken as the difference between the means of the estimated and calibrated axial position.

